# Limits to environmental masking of genetic quality in sexual signals

**DOI:** 10.1101/554477

**Authors:** James Malcolm Howie, Harry Alexander Cordeaux Dawson, Andrew Pomiankowski, Kevin Fowler

## Abstract

There is considerable debate over the value of male sexual ornaments as signals of genetic quality. Studies alternately report that environmental variation enhances or diminishes the genetic signal, or leads to crossover where genotypes perform well in one environment but poorly in another. A unified understanding is lacking. We conduct the first experimental test examining the dual effects of distinct low and high genetic quality (inbred versus crossed parental lines) and low, through high, to extreme environmental stress (larval diets) on a condition-dependent male ornament. We find that differences in genetic quality signalled by the ornament (male eyespan in *Diasemopsis meigenii* stalk-eyed flies) become visible and are amplified under high stress but are overwhelmed in extreme stress environments. Variance among distinct genetic lines increases with environmental stress in both genetic quality classes, but at a slower rate in high quality outcrossed flies. Individual genetic lines generally maintain their ranks across environments, except among high quality lines under low stress conditions, where low genetic variance precludes differentiation between ranks. Our results provide a conceptual advance, demonstrating a unified pattern for how genetic and environmental quality interact. They show when environmental conditions lead to the amplification of differences in signals of genetic quality and thereby enhance the potential indirect genetic benefits gained by female mate choice.

## 1. Introduction

Many exaggerated male sexual ornaments are thought to have evolved to provide information about the genetic quality of the signaller [1,2,3,4,5,6,7]. Yet these traits typically also respond strongly to environmental variation [8,9,10,11], and it is unclear what impact this has on their signalling function [12,13]. Does increasing environmental stress expose the underlying genetic differences in quality or mask them by overwhelming the genetic signal? Different studies have variously reported that environmental variation enhances [1,2,14,15] or diminishes [16,17,18] the phenotypic signal of genetic quality. Others reveal crossover, where genotypes that perform well in one environment do poorly in another [17,19,20,21]. These contrasting outcomes arise from a lack of consistency in experimental approach. The main problem is that analysis has focussed on genetic variation rather than distinct classes of genetic quality, coupled to a limited rather than wide range of environmental stress. We present a novel experimental design that addresses both of these deficits, which leads us to propose a unified understanding of how variation in genetic quality is impacted by environmental variation. This gives a far clearer understanding of the conditions under which sexual display traits can function to accurately reveal the genetic quality of signallers [22].

In this study, we adopt an integrated experimental approach, and for the first time examine the impact of a similar wide range in both genetic and environmental quality. We chose to focus specifically on male eyespan variation in the stalk-eyed fly [23,24] as this trait has been subject to extensive previous work. Male eyespan is highly exaggerated due to female choice [11,25,26,27,28,29,30,31,32] and also functions as a signal in male-male antagonistic interactions over mate mating sites [25,33,34]. It is highly condition-dependent relative to other traits in relation to both genetic [24, but see 36] and environmental [9,11,35] stressors, and is responsive to a range of environmental stress types [9,11,24,35,37,38], while genetically distinct families have also been shown to respond differently to environmental stress [1].

Our novel experimental design to study genetic quality-by-environment (G x E) interactions in signalling traits exploits pre-defined genetic [39,40,41] and environmental quality classes. To vary genetic quality, crosses were made within or between a set of parental inbred lines (*f* ∼ 0.908 [42]; Figure 1). This allowed us to compare low genetic quality, highly homozygous “incross” lines (*n* = 16, crosses = 67) with high genetic quality, highly heterozygous “outcross” lines (*n* = 17, crosses = 50). We used incross and outcross lines not to study the effect of inbreeding *per se*, but because previous work unambiguously shows they correspond to low and high genetic quality classes respectively [24]. The large number of independent crosses within or between lines allows us to capture the contribution of genetic variation in the sexual ornament among low and high genetic quality classes.

**Fig. 1.**
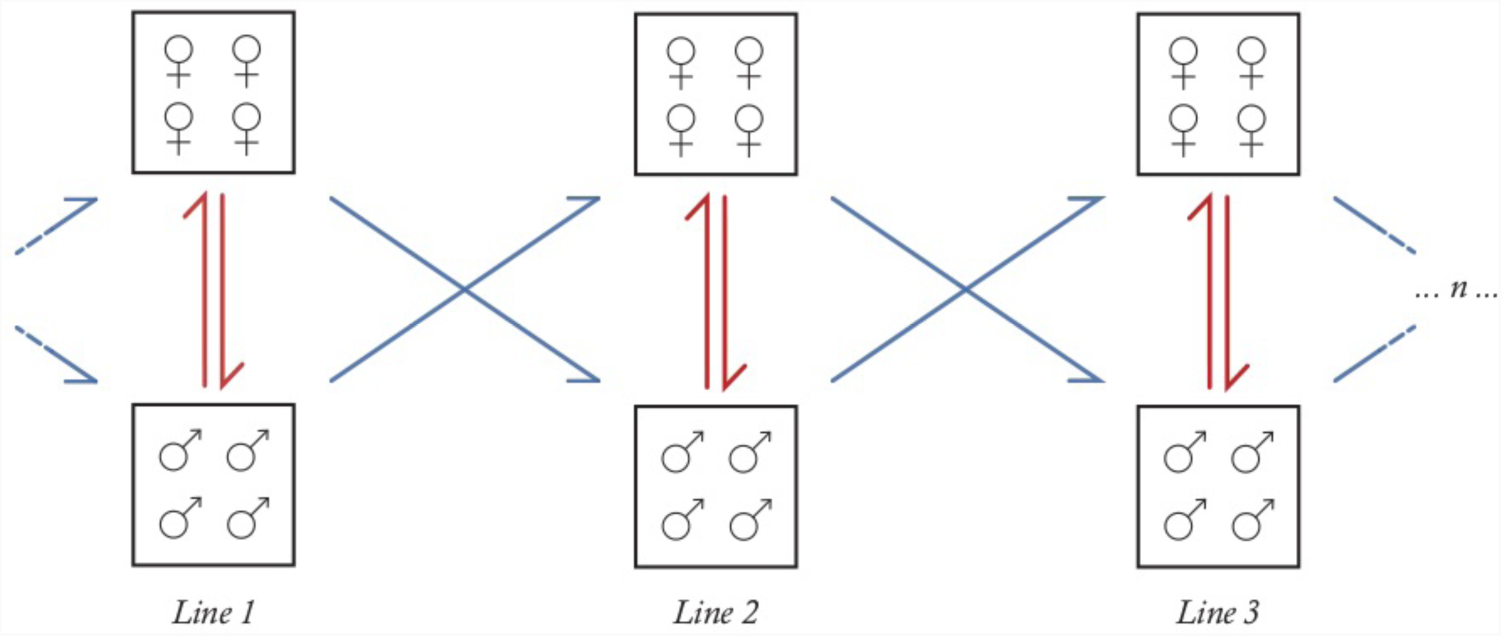
The crossing protocol used to generate incross and outcross offspring. Each inbred line was crossed with itself to create incross flies (red), or with another inbred line to create outcross flies (blue). Each cross used 4 males and 4 females and was repeated to generate two families per cross.

We likewise generated a wide range of environmental quality variation through reductions in the amount of food available to developing larvae. This approach is a well-established method for creating stress in holometabolous insects [1,11,15,26,30,35,37,40,43,44,45] and has been used extensively in prior stalk-eyed fly studies [9,11,24,30], where it generates body size variation equivalent to the range found in natural populations [32,35]. Eggs were collected from each cross, and reared under conditions of low, high and extreme environmental stress. Intermediate levels of stress between low and high were omitted due logistical constraints (the study analysed 1185 flies from 117 experimental crosses), and because those stress levels have been extensively investigated previously [11]. The extreme level was defined as the least food level where larval viability was not seriously impaired (see below)

Our use of the terminology low/high genetic quality and low/high/extreme environmental quality is necessarily arbitrary but justified in terms of the experimental design and in the results that follow. We comment further on these definitions in the Discussion. The innovation in our experimental design lies in delivering controlled manipulation of genetic quality *and* environmental stress, over several pre-defined quality levels, thereby enabling an in depth exploration of genetic quality-by-environment (G x E) variation in a male sexual ornament. We use this to investigate the signalling utility of the male ornament in providing information about indirect genetic benefits through female mate choice.

## 2. Material and methods

### (a) Production of experimental flies

A suite of 105 inbred lines were founded from a stock of *Diasemopsis meigenii* [24], an African stalk-eyed fly, derived from flies collected in South Africa by S. Hilger in 2000. After 11 generations of full-sib mating, the extant lines were bred in cage culture (∼200 flies/cage). For this study, 17 lines that varied between generations F_24_–F_33_ were chosen as parental lines to use in crosses, with flies collected as eggs laid on petri dishes containing excess puréed sweet corn (10g). At eclosion, flies were placed in large cages (15l), separated by sex (∼2 weeks) and raised until sexual maturity (∼10 weeks). Adults were fed puréed sweet corn with antifungal Nipagin, replenished twice per week. All flies were maintained using our standard protocol at 25°C on a 12:12 hour light:dark cycle, with fifteen-minute artificial “dawn” and “dusk” periods (reduced illumination) at the start and end of the light phase throughout the experiment.

### (b) Variation in genetic and environmental effects

Variation in genetic quality was achieved using previously created inbred lines [24] in a crossing protocol (Figure 1; modified from [36] and [24]). “Incross” flies were created from male-female crosses within an inbred line and “outcross” flies were created from crosses between different inbred lines. In each cross, 4 adult males from line *x* and 4 adult females from line *y* were allowed to mate in a 1.5l pot (*x* = *y* for incross, *x* ≠ *y* for outcross). Reciprocal male *x*-female *y* and female *x*-male *y* pots were set up. Multiple replicates of each cross (between 1-8) were set up, with higher rates for inbred crosses as they were less fecund. Eggs were collected twice weekly over 23 days. In all, 142 crosses were set up, of which 117 generated sufficient offspring across the food treatments: 67 incrosses of 15 inbred lines and 50 outcrosses between 16 pairs of inbred lines. An inbred line was used in an incross the same number of times as it was used in an outcross, and as far as possible equal numbers of live adult males and females were collected from each line, to balance sex chromosomal, cytoplasmic and other male/female parental effects.

Incross flies have low genetic quality as they are highly homozygous, being derived from inbred lines created by repeated brother-sister pairings (11 generations), with an expected inbreeding coefficient of *f* ∼ 0.908 [24]. In contrast, outcross flies have high genetic quality as they are expected to be heterozygous for most of the alleles fixed in the parental inbred lines from which they are derived. Although the terms – low and high genetic quality – are arbitrary, there was evidence of substantial heterosis in a variety of traits when inbred flies were crossed, so the terms reflect the nature of these two genetic groups [24].

For each incross or outcross, fertilised eggs were placed in groups of 5 in petri dishes containing two cotton pads, 15ml water and 5ml of food medium. Three qualities of food medium were used with “pure” corn diluted with water at ratios of 1:1, 1:10, and 1:20, which we designate as “low”, “high” and “extreme” stress respectively. “Pure” corn was made by forcing puréed sweet corn kernels through a fine sieve to remove husks and provide homogeneity. Food qualities were chosen based on a pilot study, with levels of food stress used that were found to lie within normal rates of egg-adult survival (Figure 2, see SI.C). Although the terms – low, high and extreme environmental stress – are again arbitrary, they nonetheless capture particular qualities. The low food level was similar to the standard media on which larvae are raised. The high food level was used previously where it was associated with reduced size in a variety of traits [24]. The extreme food level constitutes the far end of the stress spectrum before differential survival is evident (Figure 2). In the range used (i.e. 1:1 to 1:20), egg-to-adult survival did not differ in the pilot, and was at ∼50%. We did not go beyond this level, as a serious loss of adults would have placed greater logistical difficulties in delivering the already considerable sample size in the experiment. In the main experiment, a census of pupae was additionally made as a measure of survival for each cross in each environment.

**Fig. 2.**
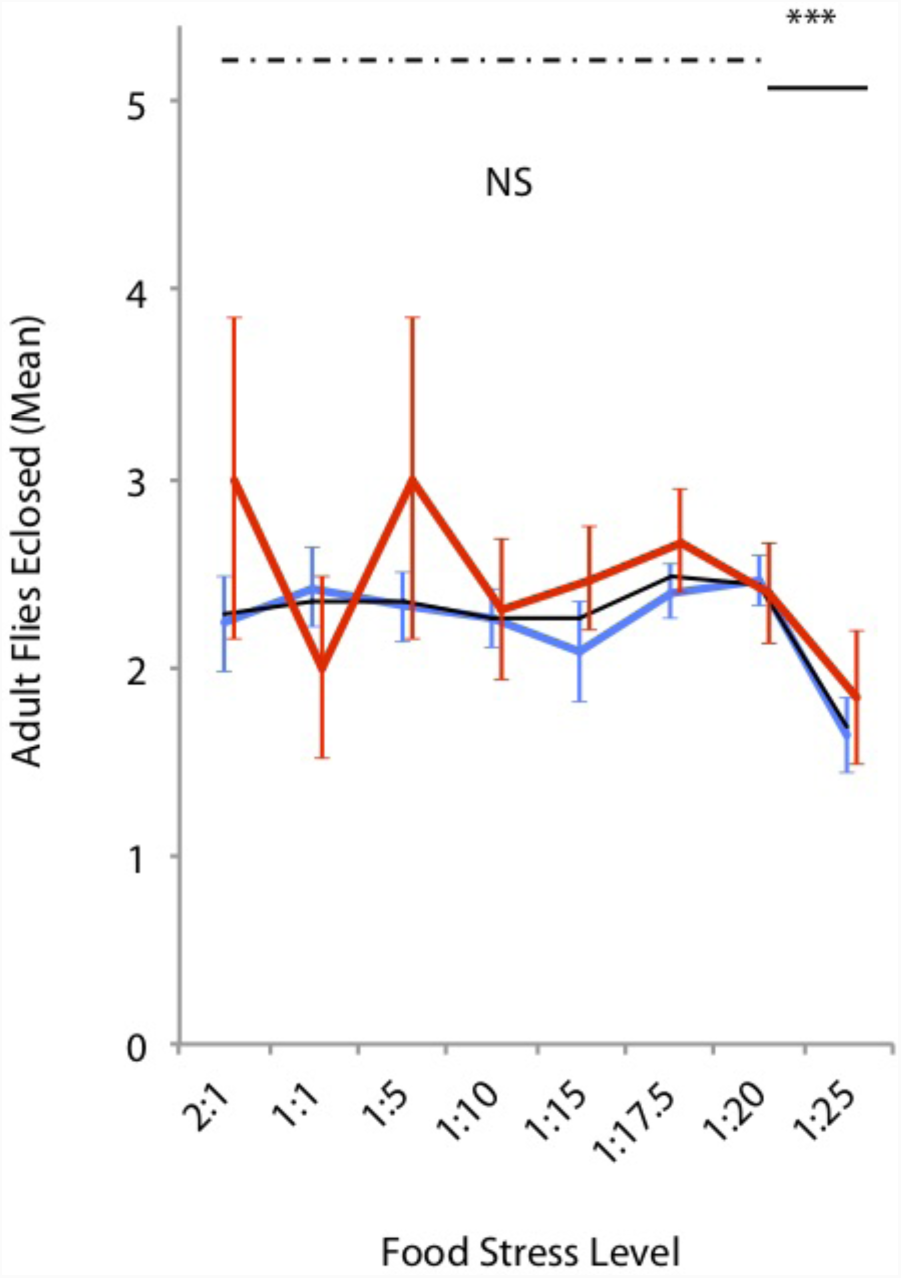
Mean number of adult flies eclosing per petri dish (± SE) given seeding with five eggs, when subject to different larval treatments (ratio of corn:water), for inbred lines (red) and stock (blue), or when pooled (black). Pairwise comparison of adjacent treatments showed a significant drop in survival between the adjacent 1:20 and 1:25 treatments (solid line, *** *P* < 0.001), and no difference between other adjacent levels (dashed line, NS). A similar pattern was observed for inbred and stock considered separately across the adjacent 1:20 and 1:25 treatments (both *P* < 0.001). Inbred and stock populations did not differ at any food level (all *P* > 0.05). Data is based on a pilot experiment (17 crosses, 10 stock, 7 inbred, *N* = 218 stock, 68 inbred; details SI.C).

### (c) Adult morphology

After eclosion, flies of each cross were collected and frozen at −20°C. All males were measured for eyespan (the distance between the outermost tips of the eyes [9,46]) and thorax (the distance between the centre of the most posterior point of the head to the joint between the meta-thoracic legs and the thorax [47,48]) to a tolerance of 0.01mm, using a video camera mounted on a monocular microscope and ImageJ image capture software v.1.46 [49]. The repeatability of these morphological trait measurements is very high at >99% [9]. In total 1186 males were phenotyped. All measurements were made blind by JMH. In a few cases (*n* = 9), a measurement was not included in the dataset due to sample damage.

### (d) Statistical analysis

To test for effects of incross/outcross genetic quality (G), environmental (E) and the G x E interaction on morphological trait variation, several general linear mixed effects models (GLMMs) were fitted via REML. In each model, G, E and their interaction were included as fixed effects. Male parental line and female parental line were included as random effects, as was cross and its interaction with E. Additional random effects of male line x E, male line x G, female line x E and female line x G explained zero variance and so were removed in model simplification. GLMMs for male eyespan had thorax added as a covariate to control for body size. Thorax length accounted for a significant portion of variance, but its addition did not substantially alter the results (for completeness, analyses of absolute trait values are given in the SI.A). GLMM models fitted pairwise to low versus high and high versus extreme environmental stress were used to further investigate the basis of the observed G x E patterns, as finally were two-tailed *t*-tests at each level of E to test whether incross male eyespan was larger or smaller than outcross male eyespan.

Coefficients of variation (CVs), the ratio of the standard deviation to the mean, were used to assess how variance in male eyespan responded to genetic quality, environmental and G x E stress. CVs control for changes in variance purely as a function of size, and are considered to be less biased than heritability estimates in G x E studies [50]. Least square means for male relative eyespan were extracted from GLMMs for each cross, for each E and G, to calculate among-cross CVs. Among-cross CVs were then compared between incross and outcross using modified signed-likelihood ratio tests (M-SLRT; [51]) in each environment, and also across environments (L-H-X), both overall and for incross and outcross. Finally, adjacent environment pairs were contrasted for among-cross CV, low with high (L-H) and high with extreme (H-X), for each genetic quality. The among-cross contrasts were conducted in the R-package ‘cvequality’ [52].

To explore the consistency of genetic lines across environments, another key aspect of G x E interactions, genetic correlations (*r*_*g*_) across adjacent environments were calculated. GLMMs were fitted with cross as a random effect and the variance component for cross was extracted for each environment. GLMMs were then carried out between pairs of adjacent environments (L-H, H-X), with the cross x E interaction included as a random effect, and the interaction variance component extracted. As before, thorax length was added as a fixed covariate to control for body size. An estimate of *r*_*g*_ was then calculated as:

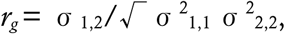

where σ^2^_1,1_ and σ^2^_2,2_ are the genetic variances in environments 1 and 2 respectively, and σ_1,2_ is the genetic covariance between the two environments [53]. Broad bounds of the *r*_g_ values were tested via model simplification and likelihood ratio tests (details in SI.A)

### (e) Statistical software used

All statistical analyses were conducted in JMP v.12.0.1 (SAS Institute 1989-2015) and R v.3.4.2 [54]. GLMM tables, effect coefficients and extended methods are shown in SI.A.

## 3. Results

### (a) Response in mean trait

As expected, male eyespan (*F*_2,45.47_ = 693.4, *P* < 0.001) and thorax (*F*_2,41.80_ = 343.4 *P* < 0.001) were smaller under higher environmental stress. The same was the case under genetic stress for eyespan (*F*_1,22.94_ = 4.783, *P* = 0.028) but not for thorax, though its response was in the same direction (*F*_1,13.78_ = 3.222, *P* = 0.095). After controlling for body size variation, the same direction of change was observed in male eyespan for environmental (*F*_2,54.66_ = 258.1, *P* < 0.001) and genetic stress (*F*_1,7.421_ = 6.203, *P* = 0.039). All following comparisons report relative trait values.

In addition, there was a genetic quality-by-environment interaction (*F*2,39.33 = 5.379, *P* = 0.009, Figure 3a). The nature of the G x E was evident from comparison of adjacent environments. The difference in male eyespan between incross flies with low genetic quality and outcross flies with high genetic quality increased from low to high environmental stress (i.e. scale variance G x E, *F*1,18.35 = 6.352, *P* = 0.021). But there was convergence between genetic quality classes after a further increase from high to extreme environmental stress (i.e. inverse scale variance G x E, *F*1,15.64 = 8.664, *P* = 0.010). This pattern was confirmed by looking at environments separately. The difference between incross and outcross male eyespan was evident at high (*t* = 8.65, df = 19.81, *P* < 0.001), but absent at low (*t* = 1.98, df = 19.79, *P* = 0.073) and extreme levels of environmental stress (*t* = −1.01, df = 21.87, *P* = 0.298).

**Fig. 3.**
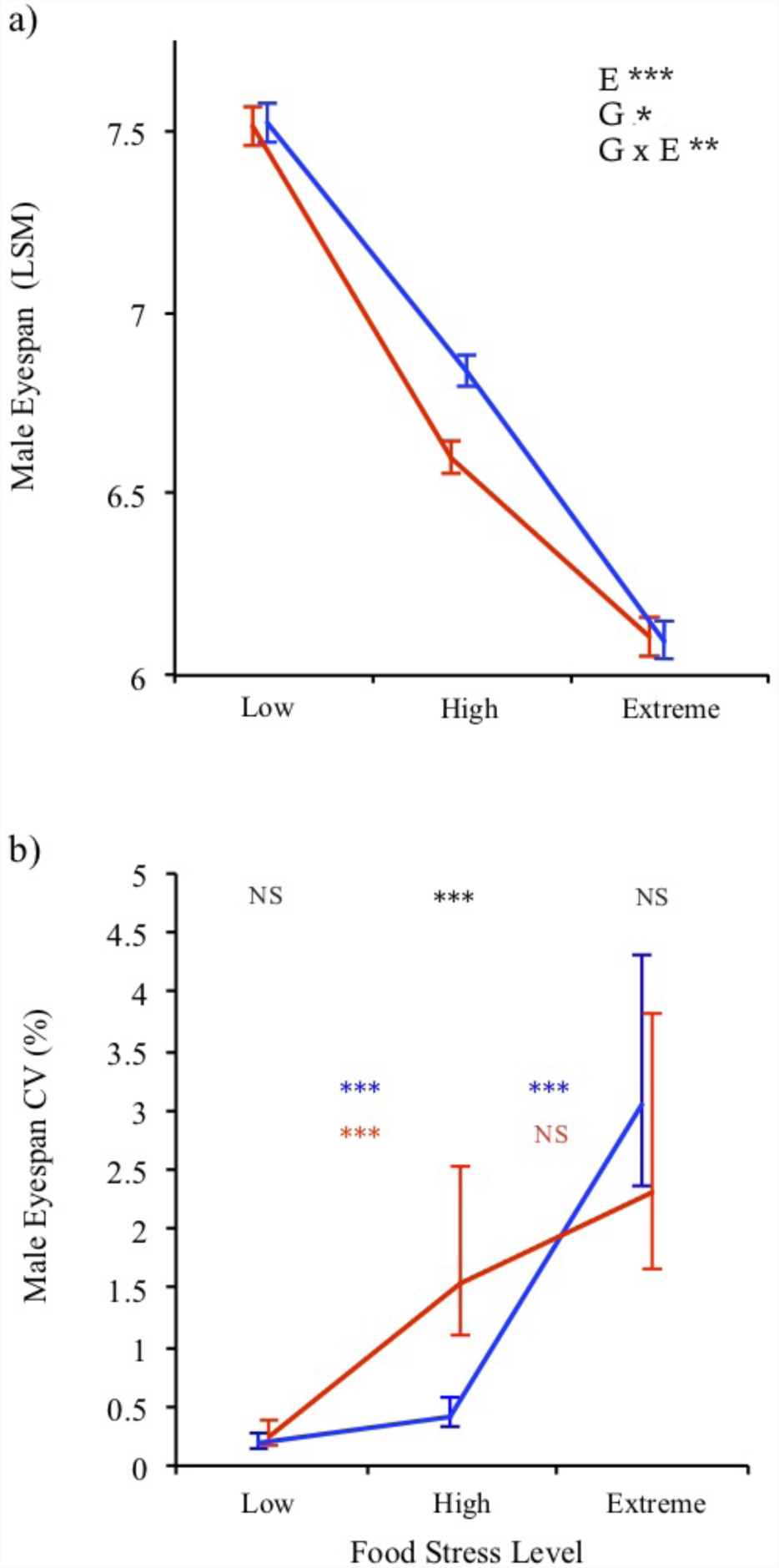
a) Male eyespan (least-squares mean ± SE) and b) coefficient of variation (CV ± 95% CI) across environmental stress (low, high and extreme) and genetic class, incross (red) and outcross (blue). The red and blue lines are shown for illustrative purposes and clarity. Asterisks denote significance: NS non-significant, * < 0.05, ** *P* < 0.01, *** *P* < 0.001. For CVs, the significance of incross versus outcross contrasts are displayed above each food level category (black asterisk at the top). The significance of within incross (red asterisks) and outcross (blue asterisks) contrasts are shown between pairs of adjacent food levels. Incross and outcross lines are jittered for clarity.

When comparisons were limited to incross lines, there were environmental 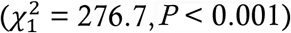 and genetic line differences 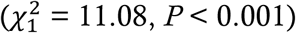 but no G x E interaction 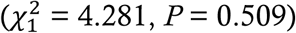. A similar pattern was found in outcross lines, where there were environmental 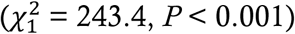 and genetic line differences 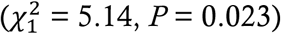 but no G x E interaction 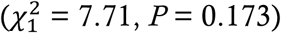. These results indicate that G x E interactions were only apparent in the comparison of genetic quality (i.e. incross vs. outcross), and not in the comparison of genetic lines within low or high genetic quality groups.

### (b) Response in trait variance

The genetic quality G x E pattern was further examined by looking at the among-cross variance in the response to stress. Coefficients of variation (CV) were used to control for the positive scaling in variance due to changes in mean trait size. Male eyespan among-cross CV (Figure 3b) was larger with greater environmental stress overall (*R*_*M*_ = 26.55, *P* < 0.001), and separately for incross (incross *R*_*M*_ = 40.00, *p* < 0.001) and outcross lines (*R*_*M*_ = 130.35, *P* < 0.001). But the extent of increase in CV from low to high environmental stress was considerably more marked among incross males with low genetic quality (1.30% increase, *R*_*M*_ = 28.95, *P* < 0.001) than outcross males with high genetic quality (0.23% increase, *R*_*M*_ = 11.95, *P* < 0.001). Differences among outcross lines were revealed to a much greater extent once the level of environmental stress increased even further, in the transition from high to extreme environmental stress (2.63% increase, *R*_*M*_ = 57.34, *P* < 0.001). This pattern contrasted again with males from incross lines, where CV did not differ between high and extreme environmental stress levels (0.78% increase, *R*_*M*_ = 1.848, *P* = 0.174, Figure 3b). As for mean eyespan, the difference between incross and outcross CV was seen only under high environmental stress (low stress *R*_*M*_ = 0.814, *P* = 0.367, high stress *R*_*M*_ = 24.32, *P* < 0.001, extreme stress *R*_*M*_ = 1.148, *P* = 0.284; Figure 3b).

### (c) Across environment genetic correlations

To further evaluate the role of male eyespan as a signal of genetic quality, we examined whether genetic lines performing well in one environment performed well across all environments (Figure 4), a critical part of the G x E pattern. When pooling all lines, there was a positive genetic correlation (*r*_g_) between low and high 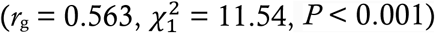, and high and extreme environmental stress 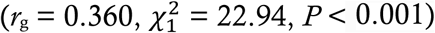. There was also a genetic correlation-by-environment interaction between low and high stress 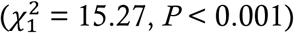 in which the genetic lines fanned out under higher environmental stress.

**Fig. 4.**
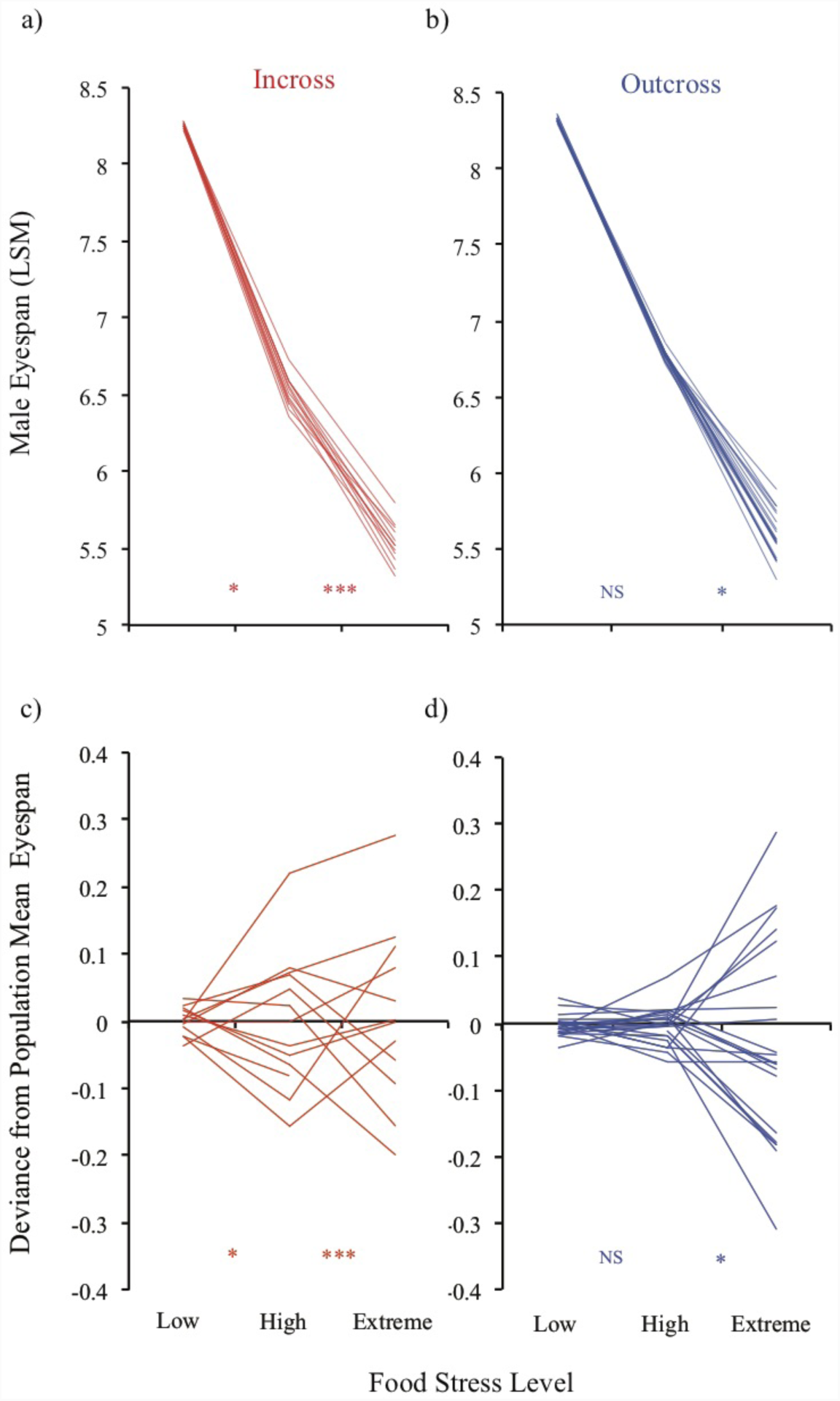
Mean male eyespan (least-squares mean relative values) at each environmental stress for each cross a) incross (red) and b) outcross lines (blue). Asterisks denote significance of the effect of cross, NS non-significant, * *P* < 0.05, *** *P* < 0.001. An alternative representation is shown as the absolute deviation of each line from the c) incross and d) outcross population mean. Error bars are excluded for clarity.

Analysing the two genetic quality classes separately, for low quality incross lines, genetic correlations (*r*_g_) were positive between low and high 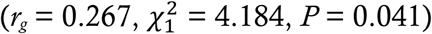, as well as between high and extreme stress environments 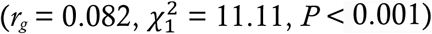. For the high quality outcross lines, there was no genetic correlation between low and high stress environments 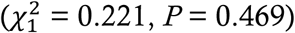, but *r*_g_ was positive between high and extreme stress environments 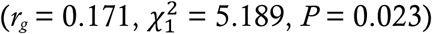. The lack of *r*_g_ was due to severely reduced variation among outcross lines in the low (CV_low_ = 0.188, CV_high_ = 0.416, CVextreme = 3.05) compared to high (*R*_*M*_ = 57.34, *P* < 0.001) or extreme stress environments (*R*_*M*_ = 88.17, *P* < 0.001; Figure 4).

### (d) Survival across G and E stress

Larval survival was measured through a census of pupae. There was a survival effect of E (*F*_1,64.36_ = 64.36, *P* < 0.001) but not of G (*F*_1,20.04_ = 0.852, *P* = 0.367) or G x E (*F*_2,64.28_ =0.976, *P* = 0.382). The effect was a reduction in survival at extreme environmental stress (pupae counts: LSM ± SE low = 2.17 ± 0.10, high = 2.29 ± 0.09, extreme = 1.55 ± 0.09). A Tukey’s HSD test confirmed that survival was lower under extreme relative to either low or high environmental stress level (*P* > 0.05). Survival did not differ between incross and outcross in any of these comparisons (all *P* > 0.05, see SI.A).

## 4. Discussion

In this study we explicitly test whether environmental stress amplifies or obscures the signal of genetic quality in male sexual ornaments. We do so in a unique way by direct manipulation of *both* genetic quality and environmental stress, the latter over multiple levels. The results enable us to put forward a unified explanation of how genetic and environmental quality interact, advancing our understanding of the genetic benefits of mate choice, with the potential to explain the diverse responses seen in other systems.

The response of male eyespan – the primary sexual ornament in *D. meigenii* – accords with previous studies in stalk-eyed flies, showing that this male ornament is a sensitive signal of both environmental [9,11,35] and genetic stress [1,24]. Of greater interest, the new data captures a full range of G x E interactions. The difference between low and high genetic quality, in both eyespan mean and variance (coefficient of variation), increases with the transition from low to high environmental food stress (Figure 3). This is an example of “scale variance” G x E in which higher environmental stress amplifies genetic differences. It has been observed across a range of species, for example in structural wing pigmentation (UV angular visibility) in the butterfly *Colias eurytheme* [55], male song attractiveness in the lesser waxmoth, *Achroia grisella* [2], and attractiveness traits in the black scavenger fly, *Sepsis punctum* [15], all examples of traits associated with sexual success. In contrast, the difference between our low and high genetic quality classes, in both eyespan mean and variance, decreases with the transition from high to extreme environmental food stress (Figure 3). This reversed pattern is an example of “inverse scale variance” G x E in which stress denudes genetic differences. It again has been observed across a range of species, for example, iridescent and orange area in the guppy *Poecilia reticulata* [17], cuticular hydrocarbon blend in *Drosophila simulans* [16], and to a more limited extent, UV brightness in *C. eurytheme* [55].

Our results are novel and striking because we see *both* scale variance and inverse scale variance in the same trait in a single species. This leads us to propose a unified hypothesis for G x E interactions in signals of quality. Moderate to large increases in environmental stress lead to amplification of the phenotypic expression of genetic quality, whereas as environmental stress becomes extreme, increases in phenotypic variation overwhelm the underlying genetic differences in quality. We note that in some previous studies, separate traits respond differently to environmental stress, suggesting variation in the threshold at which amplification transitions to restriction (e.g. [2, 55]). Future studies will be needed to identify which characteristics are associated with sensitivity levels in different traits, and whether these relate to costs of trait expression.

Yet, some evidence from other studies of sexual ornaments seems to contradict the unified hypothesis which report no interaction between genetic and environmental stress, for example in morphological traits and cuticular hydrocarbons in *D. melanogaster* [40] and several sexual traits in *P. reticulata* guppies [39]. Both of these experiments examined groups that differ predictably in genetic quality (hemiclonal lines and inbred versus outbred lines, respectively). But the lack of response likely reflects the application of insufficiently intrusive environmental stress. For example, the “stressful” environment in guppies was a moderate density [39], while that in *D. melanogaster* was a minor reduction to 70% of the normal diet [40]. A previous study in stalk-eyed flies likewise found little impact of food reduction of this order [11]. For comparison, our dilution for extreme stress was a restriction to just 5% of the standard diet. Moreover, as each of these studies used just two levels of environmental stress, analysis of complex G x E was precluded. This is not a criticism of either study, which had different goals to ours, but highlights that neither would provide an adequate test of our hypothesis. Another commonly reported pattern across diverse species, also potentially at odds with our interpretation, is “crossover” G x E in which different genetic lines are superior in different environments. This has been shown for male signal rate in the lesser waxmoth [19] and song traits of *Enchenopia* treehoppers [20]. However, “crossover” G x E is not really a distinct category, and can co-occur with “scale” or “inverse scale” G x E patterns [56]. For instance, crossover embedded within G x E scale variance patterns in the lesser waxmoth [2] and inverse scale variance in the guppy [17] has been observed. Once again, the interpretation that crossover dominates the G x E pattern requires investigation of sufficient levels of environmental stress relative to the traits in question. Without this, crossover should only be seen as part of G x E response, exerting ambiguous limits on the signalling function of the sexual ornament.

It is vitality important to examine a range of environments from low through to an extreme form of stress, alongside similar dimensions of genetic quality variation. Distinct classes of environmental quality variation were created in a standard manner through food restriction applied to developing larvae [1,11,15,30,40,45]. These treatments differed from previous studies in the use of food dilution to an “extreme”, defined as the point before larval survival showed a clear-cut decline (Figure 2). The reason for choosing this point was in part logistical, in order to easily collect similar sample size across the different stress levels. We also wanted to avoid the possibility that differential survival causes changes in trait mean and variation across the different genetic quality and environmental stresses. Despite this precaution, there was a moderate effect of the extreme environmental stress on larval survival. This could have contributed to the trait patterns observed if there was a lower level cut-off in the eyespan of survivors. We suspect this effect was minor as the mean was lowest and the CV highest in the extreme environment (Fig. 3), and more importantly, the survival deficit was equal across incross and outcross flies. Our conclusions appear to be robust. Our use of food quantity as an environmental stress was for its ease of manipulation and its use in many previous studies. Competition for food is likely to be a factor in many species and so we suspect that the results we report here are general stress responses. This needs to be established through comparison with other stresses, such as fluctuations in temperature, pH or food quality, that are part of the normal range of environmental stress in the wild [57].

To create distinct classes of genetic quality, a set of highly homozygous inbred lines (incross) were compared against crosses between lines (outcross) which are predicted to be highly heterozygous for the mutational load carried by incross lines. In the pilot experiment (Fig. 2), as well as the actual experiment, there was no difference in egg-to-adult survival between flies in the incross and outcross genetic quality treatments. The lack of a viability difference suggests that there was a strong purging of deleterious alleles during the creation of the inbred lines, as is expected and observed in other studies [58-60]. Our objective was not to study the inbreeding *per se*, as this is unlikely to be the object of female mate preference in this species. Rather we use inbreeding status as an investigative tool, in order to uncover the full nature of genetic quality-by-environment interactions on variation in signal trait size. In particular, previous G x E studies have failed to use a sufficient range of variation in genetic quality. A typical approach is to use distinct genetic lines, like brother-sister families [1,20] or inbred lines [2]. But groups that differ predictably in genetic quality have not been examined properly against a wide range of environmental stress [39-40]. Independent lines provide information about genetic variation but may differ only slightly, and unpredictably, in genetic quality, and then only with differences established *post hoc*. In our study, we distinguish between variation in genetic *quality* in the comparison of incross and outcross flies, and genetic variation between lines within these quality categories. In accordance with prior studies [1], our results show differences in performance between lines. Crucially, there was no among-line G x E once analysis was limited to a particular genetic quality class, both for incross and outcross. The set of lines in each genetic quality class appear to have been sufficiently similar in quality that they responded in an equivalent manner when challenged with our wide range of environmental stress levels (Figure 4). Only the comparison between incross and outcross flies revealed a strong G x E interaction, in which high quality (outcross) line resisted the effect of high but not extreme environmental stress.

Taking the results together allows us to comment on sexual selection on males and the potential indirect genetic benefits that arise from female mate choice. We expect sexual selection to be severely attenuated under benign and extreme environmental stress, but strong in high stress environments which amplify genetic quality differences. As stress is likely to be the norm under common ecological conditions in nature, sexual selection could often be stronger than currently estimated from laboratory experiments – typically carried out under low stress conditions of *ad libitum* food, constant temperature, no predators and parasites, and no ecological competitors. We note that our experimentation used stress from a unimodal environment variable (food availability), controlling all other physical and biotic factors, and that we used a simple measure of male signalling, leaving aside other, more subtle aspects of male behaviour used in female evaluation of their partners [46]. This implies that benign environmental conditions, equivalent to low stress in our experiment (i.e. in which larvae have excess food and little competition), are rare. Extreme environments are likewise also likely to be rare as they are not those that maintain viable populations. The majority of environments probably lie between the low and high regimes, which is consistent with the considerable range in eyespan observed among wild caught stalk-eyed flies [32].

The outcome in nature for female choice will depend on the distribution of environmental stress, its spatial and temporal variability, and hence its consequence for the pool of available mates in a given population [12]. If conditions can be categorised as low, high or extreme, then the indirect benefits of mate choice will be greatest in high stress environments, as these bring out genetic differences to the greatest extent. As genetic line correlations across environments were positive (with the exception of outcross lines between low and high food stress, where a lack of variation precluded reliable calculation), genetic differences will be evident to some extent in all environments. Where environmental conditions in a population are a mixture of low, high and extreme, individuals with the most exaggerated sexual ornaments will be an assortment of those with high genetic quality from a range of environments diluted by those less well genetically endowed but who experienced lower environmental stress during development. This cuts at the indirect genetic benefits but nonetheless there will be advantage to female mate choice. To conclude, while environmental variation places contingencies on signalling, sometimes amplifying and sometimes muting its value, genetic variation in quality between individuals will always to some extent be evident in the sexual ornament and feed through to their offspring.

## Data accessibility

Data are made available at the Dryad Digital Repository [TO ADD].

## Author contributions

JMH, KF and AP conceptualised the study and methodology, and wrote, reviewed and edited the paper. The formal analysis was carried out by JMH, who with HACD carried out the experiments. Stalk-eyed fly resources were provided by AP and KF, who secured funding and supervised the project.

## Competing interests

The authors declare no competing interests.

## Acknowledgements

The authors acknowledge support for JMH by a NERC Studentship, AP by EPSRC grants (EP/F500351/1, EP/I017909/1), and AP and KF by a NERC grant (NE/G00563X/1, NE/R010579/1). We thank Hans Feijen for sharing data from natural populations of *Diasemopsis meigenii.* Additional experimental support was provided by Rebecca Finlay, Koichi Yamanoha, and Anna Aichinger who also assisted in figure production. We acknowledge the work in creating and maintaining the inbred lines used in this work by Lawrence Bellamy, Nadine Chapman, David Ellis and Luke Lazarou.

## SUPPLEMENTAL INFORMATION

Supplemental information includes all details of statistical effect size estimates for the tests of mean effects, and additional method details. [Available after formal publication.]

